# AlphaFold2 captures conformational transitions in the voltage-gated channel superfamily

**DOI:** 10.1101/2025.03.12.642934

**Authors:** Elaine Tao, Ben Corry

## Abstract

Voltage-gated ion channels are crucial membrane proteins responsible for the electrical activity in excitable cells. These channels respond to changes in the membrane potential via conformational changes in their voltage-sensing domains (VSDs) that leads to the opening and closing of the ion conduction pore. Since alternative states of the VSDs are difficult to capture via experimental methods, we investigated the application AlphaFold2 to computationally predict structures in a range of intermediate and endpoint states. By generating 600 models for 32 members of the voltage gated cation channel superfamily we show that AlphaFold2 can predict a range of diverse structures of the VSDs that plausibly represent activated, deactivated and intermediate conformations with more diversity seen for some VSDs compared to others. Modelling the full sequence of pseudo-tetrameric channels also produced a range of diverse states in the pore and intracellular regions that potentially represent physiologically relevant states. However, there are some incongruities between certain generated models and resolved cryo-EM structures suggesting that further validation is required to confirm their structural and functional relevance.

## Introduction

The voltage-gated (VG) cation (K^+^, Na^+^ and Ca^2+^) channel superfamily consists of tetrameric or pseudo-tetrameric proteins that facilitate both selective and non-selective ion permeation across cell membranes in response to changes in membrane potential (Fig. 1A). These channels are critical for regulating the electrical signalling between excitable cells throughout the body. The overall architecture of channels in this family is conserved, with four identical or analogous domains formed by one, two or four separate protein units. Each of these domains contain voltage-sensing domains (VSDs) made up of a four-helix bundle (Fig. 1B-C), as well as, an additional two helices and a re-entrant pore loop that form the ion-conducting pore, activation gate and determine ion selectivity.

**Figure 1.**
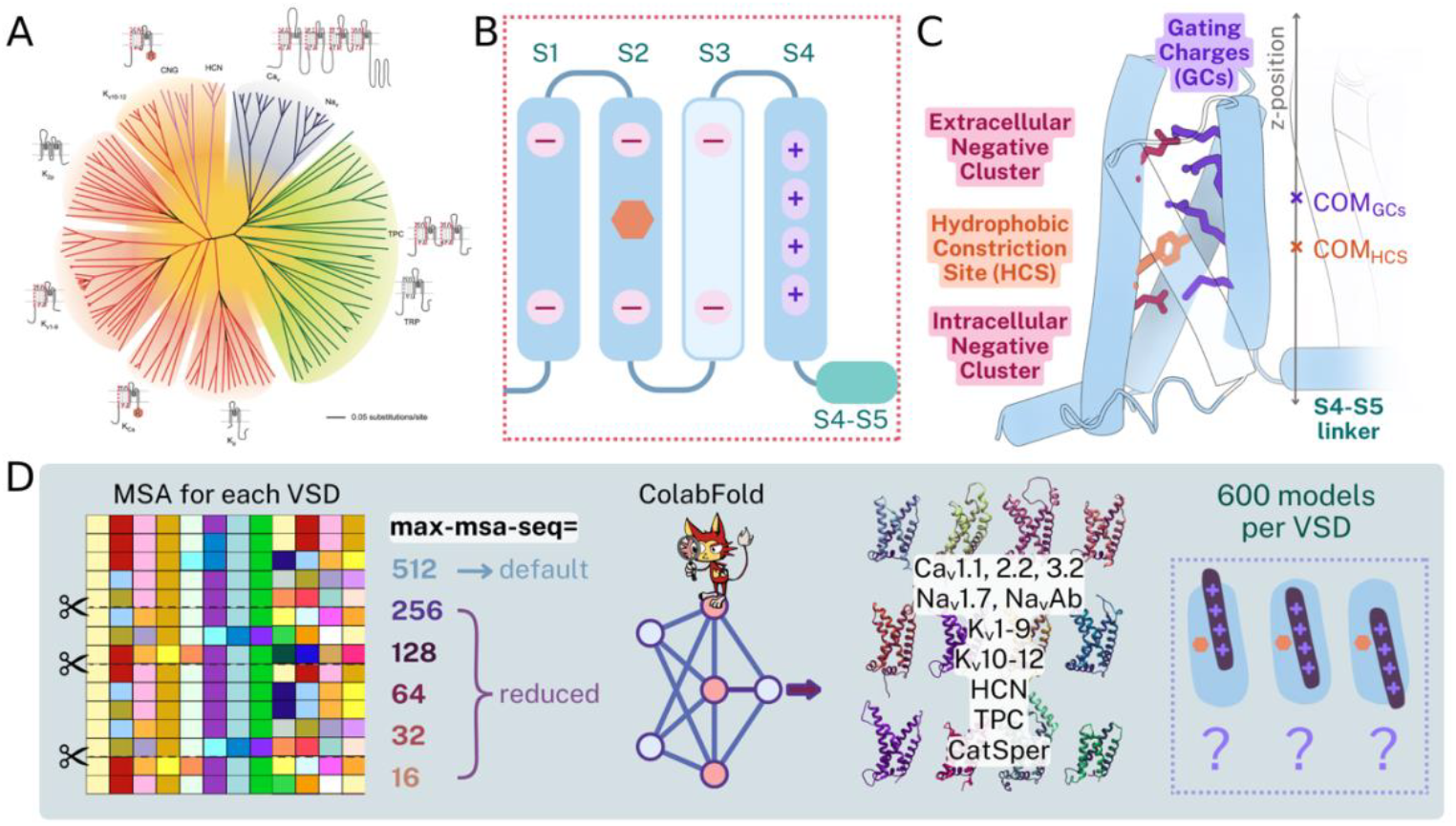
Examining VSD conformations in the VG cation channel superfamily. The VG cation channel superfamily, anatomy of a VSD and the measurement of S4 helix displacement. (A) An unrooted phylogenetic tree of the VG cation channel superfamily, adapted from Yu and Catterall (2004). (B) Topology of a typical VSD, featuring negatively charged (or sometime polar) residues located on the S1-S3 helices, the hydrophobic constriction site (HCS) at the midpoint of S2, and the four main positively charged gating residues on S4. (C) Example structure of a VSD showing the residues described in B and coloured accordingly; as well as showing how the S4 z-displacement is measured using the z-position of the COM of the four main gating charges relative to that of the HCS. (D) Schematic summarising the procedure for AF2 structure prediction with subsequent restrictions applied to depth of the MSA input.

The S4 helix, which contains the positively charged residues that facilitate the voltage-sensitive gating process (Fig. 1B-C), is one of the most scrutinised transmembrane helices in the proteome. There is a long history of experiments addressing VSD function many of which attempt to understand how the charged S4 helices translocate across the membrane. Experiments using a fluorescent probe attached to the S4 have directly tracked its movement during voltage-dependent gating for K^+^ channels (Mannuzzu et al., 1996), Na^+^ channels (Chanda and Bezanilla, 2002) and Ca^2+^ channels (Wang et al., 2024). X-ray crystallography and cryo-EM experiments resolve the default state of VSDs in the absence of a transmembrane potential, which tends to be an activated state. By applying experimental manipulations such as stabilising mutations (Huang et al., 2022), VSD-trapping toxins (Jiang et al., 2021, Wisedchaisri et al., 2021), or inducing a potential across liposomes (Mandala and MacKinnon, 2022), recent structural studies have managed to capture some alternative states of different VSDs. Molecular simulations of K^+^ channels have also been used to examine the VSD activation/deactivation transition pathway, revealing key intermediate states and secondary structural changes during the process, as well as, quantifying their energetics and kinetics (Delemotte et al., 2011, Schwaiger et al., 2011, Jensen et al., 2012).

Movement of the S4 helix is coupled to the pore domain of ion channels, which upon VSD activation opens to conduct ions across the membrane. Abundant structural studies of the various ion channels in the superfamily have resolved the transmembrane region of the pore domain comprising of the S5 and S6 helices. The pore gate is formed by the centre-facing hydrophobic residues on the intracellular end the four S6 helices that seal the pore in the non-conductive or closed state. In addition to voltage sensing and pore gating, Na^+^ channels also feature a distinct inactivation mechanism, whereby the hydrophobic isoleucine-phenylalanine-methionine (IFM) motif on the flexible DIII-IV linker binds near the pore gate to force the protein into a non-conducting conformation (Yan et al., 2017).

AlphaFold2 (AF2) is an artificial intelligence-based 3D protein structure prediction tool that was trained on experimentally resolved structures from the Protein Data Bank and evolutionary sequence data in the form of multiple sequence alignments (MSAs) (Jumper et al., 2021). A challenge for protein structure prediction, however, is that proteins do not adopt a single static structure. Instead, they have dynamic structural distributions with functional regions that fluctuate on different timescales, as is evident in the VSD of ion channels. Recent studies have used AF2 to generate structural ensembles for membrane proteins that undergo conformational changes, including receptors and transporters (del Alamo et al., 2022). To accomplish this, various strategies to mask or restructure the MSA input have been applied, such as reducing MSA depth (del Alamo et al., 2022) (Fig. 1D), and clustering the MSA (Wayment-Steele et al., 2022). Some conformational diversity is evident in AF2 models of Na_v_ and HERG channels (Lopez-Mateos et al., 2024, Ngo et al., 2024), showing that AF2 can predict diverse conformations for specific regions of the channel (i.e. pore gate, selectivity filter, voltage sensors). However, assessing whether these conformations represent real structural transitions remains challenging. Moreover, these studies have not leveraged the structural similarities across the VG cation channel superfamily to assess AF2’s reliability in generating diverse conformations.

Since it is challenging to resolve all the functional states of ion channels and transition intermediates using either experimental structure determination or MD simulations, we explored whether AF2 could offer a less resource-intensive way to probe the conformational space of VG cation channels. Here we investigated the capacity of AF2 predictions to reflect (1) localised conformational changes of the S4 helix in the VSD, and (2) broader structural heterogeneity across the entire channel tetramer, including conformational states of the pore and intracellular machinery. In the first part, we used AF2 to generate models of 32 different VSDs from representative members of the VG cation channel family. These include VSDs from the four domains of pseudo-tetrameric sodium- and calcium-selective channels, VSDs from homotetrameric potassium-selective and other cation-selective channel families, as well as VSDs from the two domains of the two-pore channels. In the second part we, we generated complete tetrameric models of two pseudo-tetrameric channels including the pore domains and intracellular regions. In both cases we assessed the structural diversity produced by AF2 after applying both the default MSA depth and a range of reduced MSA depths to enhance conformational diversity.

## Results

Within the ‘voltage-dependent channel domain superfamily’ (as classified by InterPro (Blum et al., 2025)), there are over 300 structures resolved from a variety of eukaryotic and prokaryotic species. The entries in this superfamily that are from homo sapiens includes examples from 8 of the 9 voltage-gated sodium (Na_v_) channels and the sodium leak channel NALCN, 8 of the 10 subtypes of voltage-gated calcium (Ca_v_) channels, and only 12 of the 40 known voltage-gated potassium (K_v_) channel subtypes. Interestingly, other proteins like the voltage-sensing phosphatases (VSPs) and transient receptor potential melastatin/polycystin channels (TRPM8, TRPP1, TRPP2) have also been classified in this group.

### Different degrees of conformational diversity across VSDs generated by AF2

To evaluate the conformational heterogeneity generated by AF2, we generated 600 models of each of 32 different VSD sequences. K_v_ channel examples are a single domain, whereas Ca_v_ and Na_v_ channels each includes four domains of different protein sequences (shown as D1-D4). Although the two-pore channel TPC1 contains two VSD-like regions, only the first was identified in the InterPro database.

Across the VSD family, there is significant variation in the VSD models produced by AF2 that are distributed over a spectrum of activated (up) and deactivated (down) conformations (Fig. 2). The distributions include structures predicted using default MSA parameters (100 models, in blue), as well as five levels of MSA depth reduction (500 models combined, in maroon). The z-position of the S4 helix relative to the HSC (Fig. 1C) was used as the measure of VSD conformation. A zero value on the displacement scale represents conformations where the S4 midpoint is in line with the HCS. Overall, we see that reducing the MSA depth leads to a broader distribution of states, as well as shifting it toward the ‘down’ VSD conformation.

**Figure 2.**
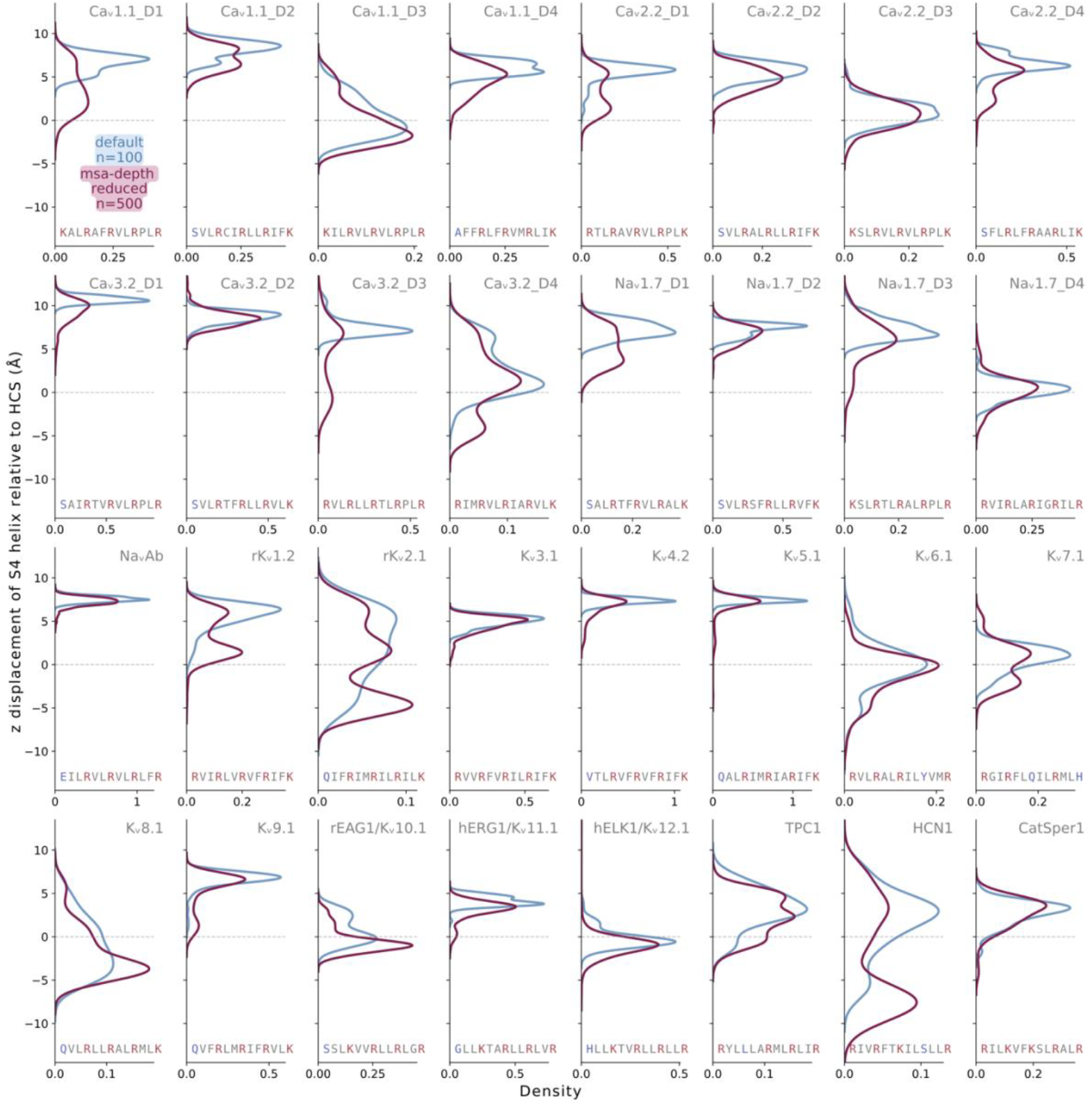
Predicted VSD models display conformational diversity. The z displacement of the S4 helix across 600 AF-generated models for 36 different ion channel VSD sequences. VSD models were aligned to their embedded positions in the membrane with zero denoting the approximate z-position of the HCS. Sequences of the S4 helix are shown for each VSD, spanning the region between the four gating charges (canonically known as R1, R2, R3 and R4 from Na_v_Ab) used for measurement plus the three residues preceding this stretch (as some VSDs also have a positively charged residue in this region).

We group the distributions of VSD position seen for each channel (Fig. 2) into three scenarios. (1) A homogenous distribution of conformations with models generated with S4 in the up position. In these, the distribution peaks are very narrow and centred at values around or greater than 5 Å – for example Ca_v_3.2-D1 and -D2, Na_v_1.7-D2, Na_v_Ab, K_v_3.1, K_v_4.2, and K_v_5.1. (2) A distribution showing more conformational diversity in the displacement of the S4 helix with a peak number of models generated in an intermediate position with the S4 midpoint level with the HCS – for example Ca_v_1.1-D3, Ca_v_2.2-D3, Na_v_1.4-D4, EAG1 and ELK1). (3) Multi-modal distributions covering an extensive range of S4 displacements, suggesting greater conformational heterogeneity and a variety of intermediate VSD states. In some of these cases, the distributions are bimodal, suggesting two distinct stable states – one located around the 0 Å mark and another one in the up state where the S4 is ∼6 Å above the HSC. This is clearly seen for Ca_v_1.1-D1, Ca_v_2.2-D1, Ca_v_3.2-D3, K_v_1.2, K_v_9.1. In other cases, there are three peaks – the two identified previously, plus an additional peak in the down state where S4 is ∼5 Å below the HCS. This is evident for Ca_v_3.2-D4 and K_v_2.1. In general, we see that MSA depth reduction biases the distributions towards more down-state VSD models with a more pronounced peak in the lower range.

To better understand the variety of VSD structures represented by these distributions, we classify them according to how many gating charges are found below the HCS. As seen in Fig. 3A, AF2 can predict structures of nearly all VSDs in more than one of these states. Next, we closely examine four representative examples from the different VGCC families with the greatest conformational diversity – Ca_v_3.2-D4, Na_v_1.7-D4, K_v_2.1 and HCN1 (denoted with * in Fig. 3A). For K_v_2.1 the three peaks each correspond to the repositioning of one gating charge (Fig. 3B); that is, the topmost peak represents the state where one gating charge falls below the HCS (pink), the middle peak represents two gating charges below the HCS (blue) and the bottom peak represents three gating charges below the HCS. A similar range of conformations is observed for Ca_v_3.2-D4, however the middle peak is the most frequently observed with two gating charges below the HCS (blue). In the case of Na_v_1.7-D4, the distribution of VSD states is even more skewed towards the middle, almost appearing to be a singular peak, but with a small number of structures found in the other two states. For these three cases, the distributions represent the movement of two gating charges past the HCS, giving rise to three peaks in the S4 z displacement distributions. Conversely, the VSDs presenting with bimodal distributions represent the movement of one gating charge.

**Figure 3.**
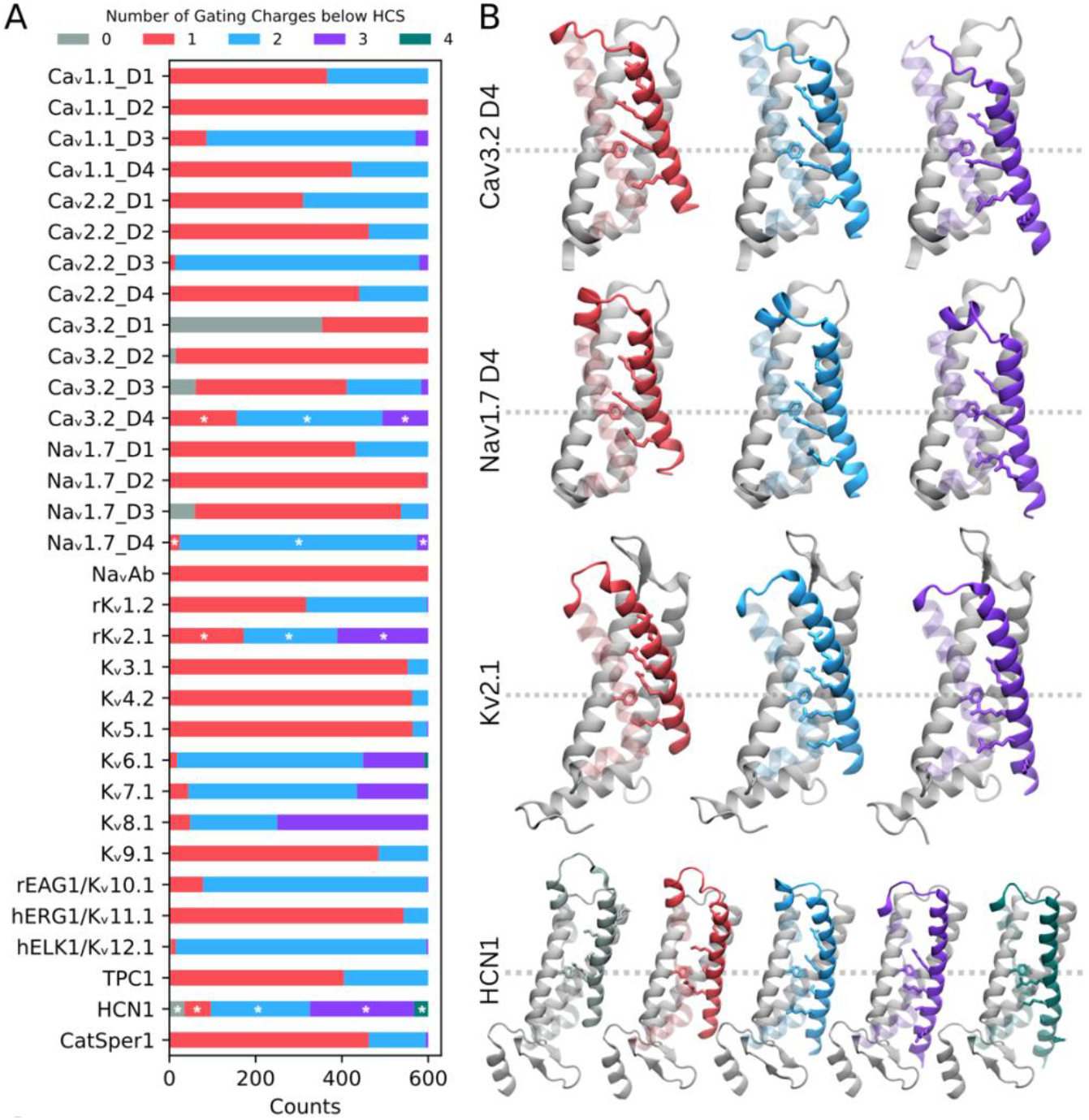
Distinct conformational states sampled by AF2 predicted VSD models. (A) The count of VSD models with zero, one, two, three, or four gating charges below the HCS determined for all 600 AF2 models for 36 different VSD sequences. (B) Representative snapshots of different gating charge conformations from different examples of Na_v_, Ca_v_, K_v_ and nucleotide-gated (HCN) channels. The colours and models shown are reflect the colour and location of the * seen in panel A.

**Figure 4.**
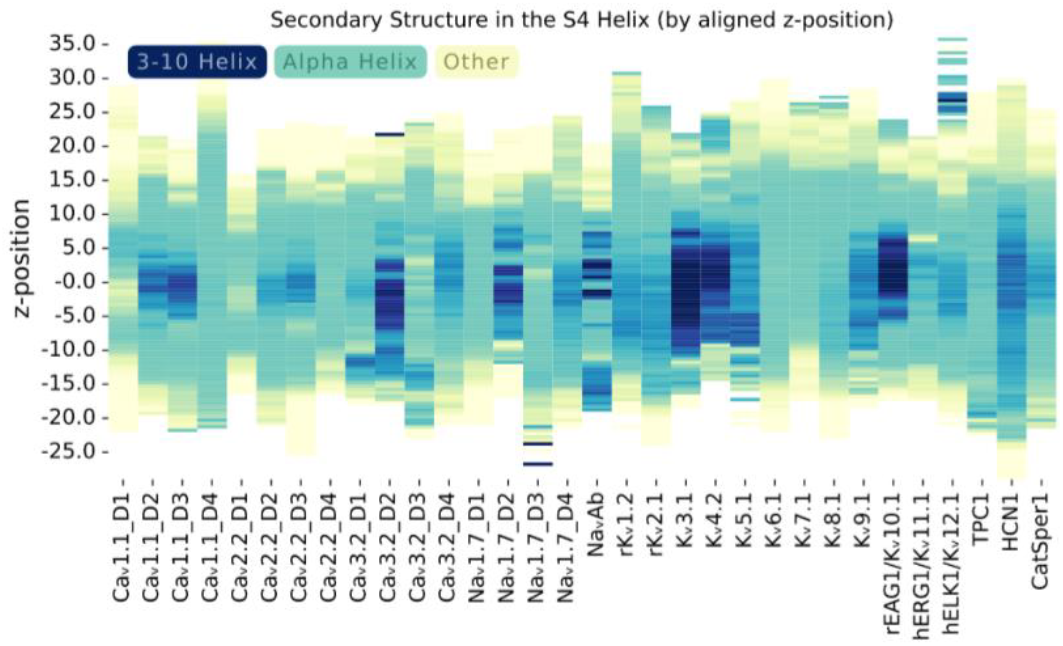
AF2 predicted models show secondary structure variation within the S4 helix. The presence of 3-10 helical regions across the S4 helices is plotted against z-position for all the predicted VSD models.

The VSD from HCN1 exhibits the broadest spectrum of states, spanning a displacement of over 20 Å and grouped into two distinct peaks – one peak where the midpoint of the S4 gating charges is above the HCS and the other where the S4 midpoint falls below (Fig. 2). However, we observe that the ‘up’ population represents three possible stable conformations (Fig. 3) – no gating charges (grey), one gating charge (pink) and two gating charges (blue) below the HCS, whereas the ‘down’ population represents two states – three gating charges (purple) and four gating charges (green) below.

To further assess the AF2-generated VSD structures, we analyse the secondary structure of the S4 helix. Previous work has established the necessity to form a short 3-10 helical section within S4 to facilitate the sliding helix transformation during VSD activation and deactivation of K_v_ channels (Henrion et al., 2012). Across the different channel VSDs, we observe differences in the proportion of 3-10 helix in the S4 of the AF2 models. Some VSDs have S4 helices that are predominantly alpha helical in nature (i.e. Ca_v_1.1-D4, Ca_v_2.2-D4, Na_v_1.7-D1, K_v_6.1, K_v_7.1, K_v_8.1). In contrast, K_v_3.1 features the most prominent 3-10 helical region. Since conformations of the K_v_3.1 VSD are fairly homogenous, the 3-10 helical structure spans almost the entire length of the helix. In contrast, given the diversity in S4 positions predicted for HCN1, the z-position of the 3-10 helix shifts depending on the state of the VSD. Thus, this highlights the diversity in S4 secondary structure across the VSD superfamily and differences in their structural dynamics during the process of activation/deactivation. Further work is needed to validate these observations.

### AF2-generated VSD conformations are sensitive to variations in the input sequence

In our tests of AlphaFold, we observed that changing certain parameters also caused a shift in conformational distributions in the VSDs. For example, the addition of the S4-S5 linker (a stretch of 14 residues) preceding the S4 led to predictions of conformational distributions that were biased towards the up/activated state. This was observed for both K_v_2.1 and Na_v_1.7-DIV (Fig. S2).

To determine whether the conformational heterogeneity was impacted by the presence of the pore domain in the case of a complete ion channel, we generated models of the full pseudo-tetrameric sequences for the Na_v_ channel (Na_v_1.7) and a representative Ca_v_ channel (the T-type Ca_v_3.2 channel) applying the same parameters and MSA depth restrictions as above. Structural predictions made with the full channel sequence, display different distributions of VSD conformations compared to models made using the VSD sequence alone (Fig. 5 A-B). For most VSDs, the addition of the pore domain leads to a downwards shift in the distributions.

**Figure 5.**
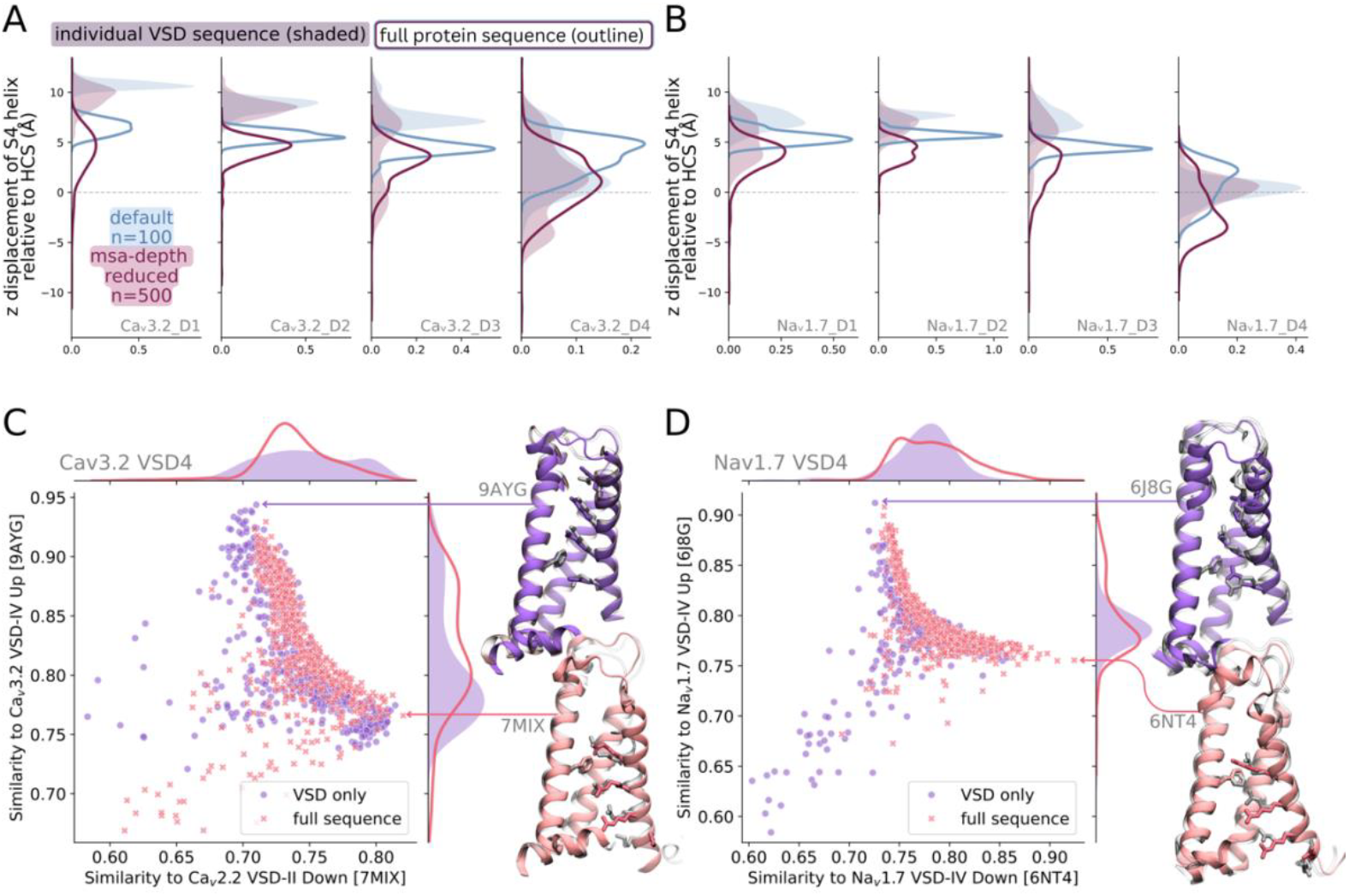
VSD conformational distributions show slight differences when predicted alone or together with the rest of the protein but resemble resolved structures. (A-B) Distributions of the S4 helix z-displacement measured across VSD only AF2-generated structures (shaded) and full tetrameric sequences (outline) for Ca_v_3.2 (A) and Na_v_1.7 (B). (C-D) TM score similarity between AF2-generated models of VSD4 compared to resolved cryo-EM structures of VSDs in up and down states for Ca_v_3.2-VSD4 (C) and Na1.7-VSD4 (D).

To assess conformational diversity in the VSD models in the absence of intracellular regions, we first generated models of Na_v_1.7 with the intracellular domain linkers replaced with chain breaks using AF2-multimer (Fig. S3, top row). This resulted in less conformational heterogeneity than seen in AF2 models of the full sequence (second row), full sequence with intracellular regions replaced with glycine chains (third row), or individual VSDs (bottom row). The distributions were also not significantly altered by reductions in MSA depth.

Interestingly, the fourth VSDs of both Na_v_1.7 and Ca_v_3.2 exhibited the greatest range of conformations. To assess the feasibility of the AF2-generated conformations of these two VSDs, we compared them to existing resolved structures using the metric of TM score (Fig. 5 C-D). The AF2-generated Ca_v_3.2-VSD4 model that was most similar to a resolved structure had a TM score of ∼0.95 compared to the cryo-EM structure of the up conformation, representing a very high degree of similarity. This can also be seen in the aligned AF2 and cryo-EM structures (Fig. 5C, AF2 model in purple, Ca_v_3.2 VSD-IV cryo-EM structure in transparent gray). As there are no resolved down states of Ca_v_3.2-VSD4, we compared to the resolved down state of Ca_v_2.2-VSD2. In this case, the TM score of the most similar model was slightly lower (∼0.82) compared to that seen for the up state, which may be partly due to the comparison being made to a different domain and subtype. However, there is still strong overall agreement, with three of the gating charges aligning well despite some structural deviation at the top of the S4 helix (Fig. 5C, AF2 model in pink, Ca_v_2.2 VSD-II cryo-EM structure in transparent gray). For Na_v_1.7-VSD4, the most similar models to resolved cryo-EM structures of the up/down states all have a TM score >0.9 (Fig. 5D), showing that AF2 can predict conformations that closely resemble the distinct activated and deactivated VSDs of Na_v_1.7 in cryo-EM structures.

For these two cases, we have shown in detail that the AF2-generated models not only produce up and down conformations of the VSD, but also a distribution of structures that provides a clear pathway between the two endpoints (Fig. 5 C-D). Therefore, structures at the leading edge of these clusters likely represent plausible transitional states, offering insight into the mechanistic process of VSD activation.

However, in contrast, the HCN1 models may not as accurately represent the native structures as seen for Na_v_1.7 or Ca_v_2.2. HCN1 contains a long S4 helix with two additional charges below the four measured ones. Compared to resolved structures of the activated S4 state, the topmost state of the AF2 models is misaligned by approximately one helical turn below the corresponding cryo-EM structure (Fig. S4). Conversely, the range of modelled down states extend past what is seen in the cryo-EM structures of the deactivated HCN1 VSD (Fig. S4). The resolved cryo-EM structure of the down conformation also features a bend in the S4 helix that is not present in the AF2 models. This suggests the conformational diversity in HCN1 models are not as well representative of physiological states.

### Other conformational changes observed for the complete channel tetramers

Across predicted structures of the complete pseudo-tetrameric Ca_v_3.2 and Na_v_1.7 channels, we also observe a diversity of conformations within the pore domain. To ensure that analysis did not include poorly folded models, we checked the accuracy of the overall folding by measuring the TM score of the Na_v_1.7 AF2 models to the resolved cryo-EM structure 6J8G (Fig. S5). All 100 structures in the default MSA and the first three MSA depth restrictions (max-msa-seq = 256, 128, 64) have a TM score > 0.9, indicating a correctly folded state. However, if max-msa-seq is reduced to 32 and 16, the percentage of incorrectly folded models increases. Interestingly, the S4 helix is predicted on average with a lower pLDDT compared to the other five transmembrane helices, likely due to its position changing the most between the functional states of the channel (Fig. S5). For further analysis of conformational states, models below a TM score of 0.85 are discarded.

Significant conformational heterogeneity is observed in the pore for the models of Ca_v_3.2 and Na_v_1.7, which is quantified by the distances measured between the COM of opposing pore gate residues (Fig. 6A-C). To differentiate between key conformations, the graph is divided into quadrants – where the lower left represents the most narrow or closed pore gate, the upper right the most open pore, and the remaining two represent asymmetric states (Fig. 6A-B). Example structures of each of these conformations are shown for Na_v_1.7 (Fig. 6C). Overall, reducing the MSA depth leads to a broader distribution of distances and pore conformations. Na_v_1.7 models show a bias towards the asymmetric conformation where the D1 and D3 helices are closer together than the D2 and D4. Notably, the pore models become increasingly biased towards this state with subsequent reductions in the MSA (Fig. 6B). This asymmetry is less apparent for Ca_v_3.2 and there is no obvious shift in the distributions with MSA reduction (Fig. 6A).

**Figure 6.**
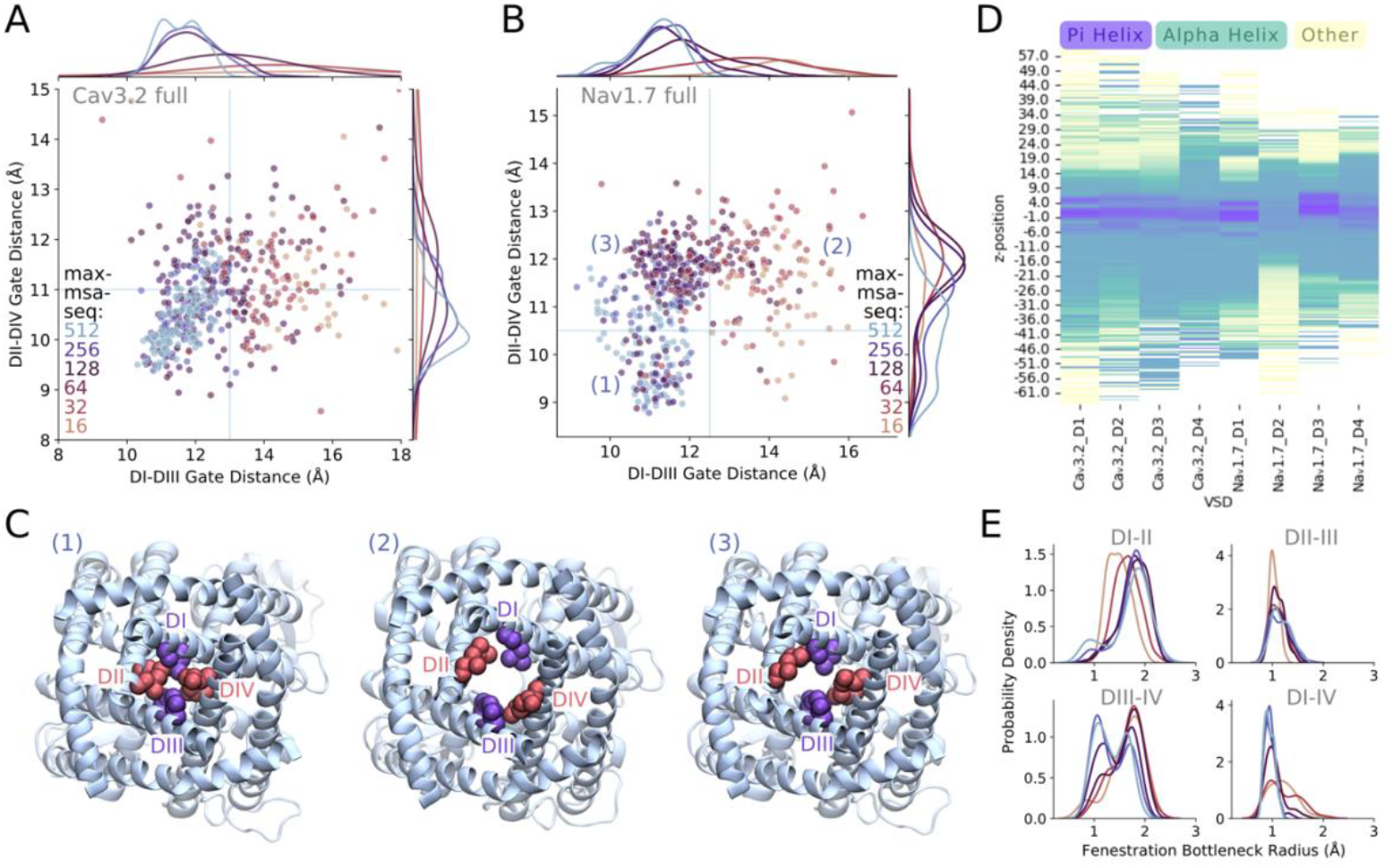
Conformational heterogeneity in pore gate and S6 helices measured in predictions of the full tetrameric sequences of Ca_v_3.2 and Na_v_1.7. (A) Scatterplot showing the centre-of-mass (COM) distances between two hydrophobic residues located on opposing domains of the Ca_v_3.2 pore gate; residues used for COM calculations for each of the domains were L415 and A419 on DI, V1011 and V1015 on DII, V1551 and V1555 on DIII, and V1856 and M1860 on DIV; points coloured by level of MSA depth restriction applied during AF2 predictions. (B) Scatterplot showing the COM distances between two hydrophobic residues located on opposing domains of the Na_v_1.7 pore gate; L398 and A402 on DI, L964 and L968 on DII, I1453 and I1457 on DIII, and I1756 and L1760 on DIV; points coloured by level of MSA depth restriction applied during AF2 predictions. (C) Representative structures of the Na_v_1.7 pore from three of the quadrants indicated in B, where 1, 2 and 3 represent a closed, open and asymmetric pore gate, respectively; gate residues are shown in Van der Waals representation. (D) Distributions of bottleneck radius across the four fenestrations of Na_v_1.7 AF2-generated models; distributions coloured by level of MSA depth restriction similar to in A and B.

In addition to opposing domain pore gate distances, we also analysed fluctuations in the secondary structure of S6, as pi-helical regions are frequently reported at the midpoint of this helix in cryo-EM structures and examined in simulations eukaryotic Na_v_ channels (Choudhury and Delemotte, 2023). Our AF2-generated models recapitulate this, exhibiting a hotspot for pi-bulges between z-positions of -5 Å to 5 Å at the midpoint of the S6 helices (Fig. 6D). In Ca_v_3.2, this the pi-bulge also appears is uniformly across the S6 of all domains, whereas in Na_v_1.7 the pi-bulge is apparent in the S6 D1 and D3, to a lesser extent in D4 and hardly at all in D2.

For the Na_v_1.7 models, there is variation in the sizes of the four membrane-facing fenestrations, located between the domains of the pore (Fig. 6E). The bottleneck radius measures the narrowest point along each fenestration. The fenestrations DII-III and DI-IV are typically quite narrow and restricted across all AF2-generated models, with their bottleneck radius distributions centred around 1 Å. In contrast, DI-II and DIII-IV have a wider conformation – a second peak in bottleneck radius distribution seen ∼1.8 Å. Interestingly, with increasing MSA depth restrictions, fenestration bottleneck radii become more skewed towards the wider state.

To evaluate conformational diversity in the inactivation gate of Na_v_1.7, we measured the distance between the IFM motif and its receptor site, and the distance between conserved residues on the C-terminal end of DIV-S6 and the DIII-IV linker involved in ‘switch 2’ (Clairfeuille et al., 2019), E1769 and K1487 respectively. Across the AF2-generated ensembles of the full Na_v_1.7 sequence (Fig. 7), we observe two distinct populations – (1) structures with the IFM motif associated with the receptor site wedged near the pore gate, and the DIII-IV linker orientated with K1487 facing away from the pore; (2) structures with the IFM motif dissociated >20 Å away from its receptor and K1487 interacting with E1769 on the C-terminal region. There is a lack of feasible structures in the transitional zone between the two states, unlike what is seen for the VSD ensembles. Additionally, there is no obvious correlation between MSA depth restriction and the likelihood of generating one conformation or the other. Thus, the default application of AF2 can reproduce the same variability of states, potentially representing physiologically relevant conformations.

**Figure 7.**
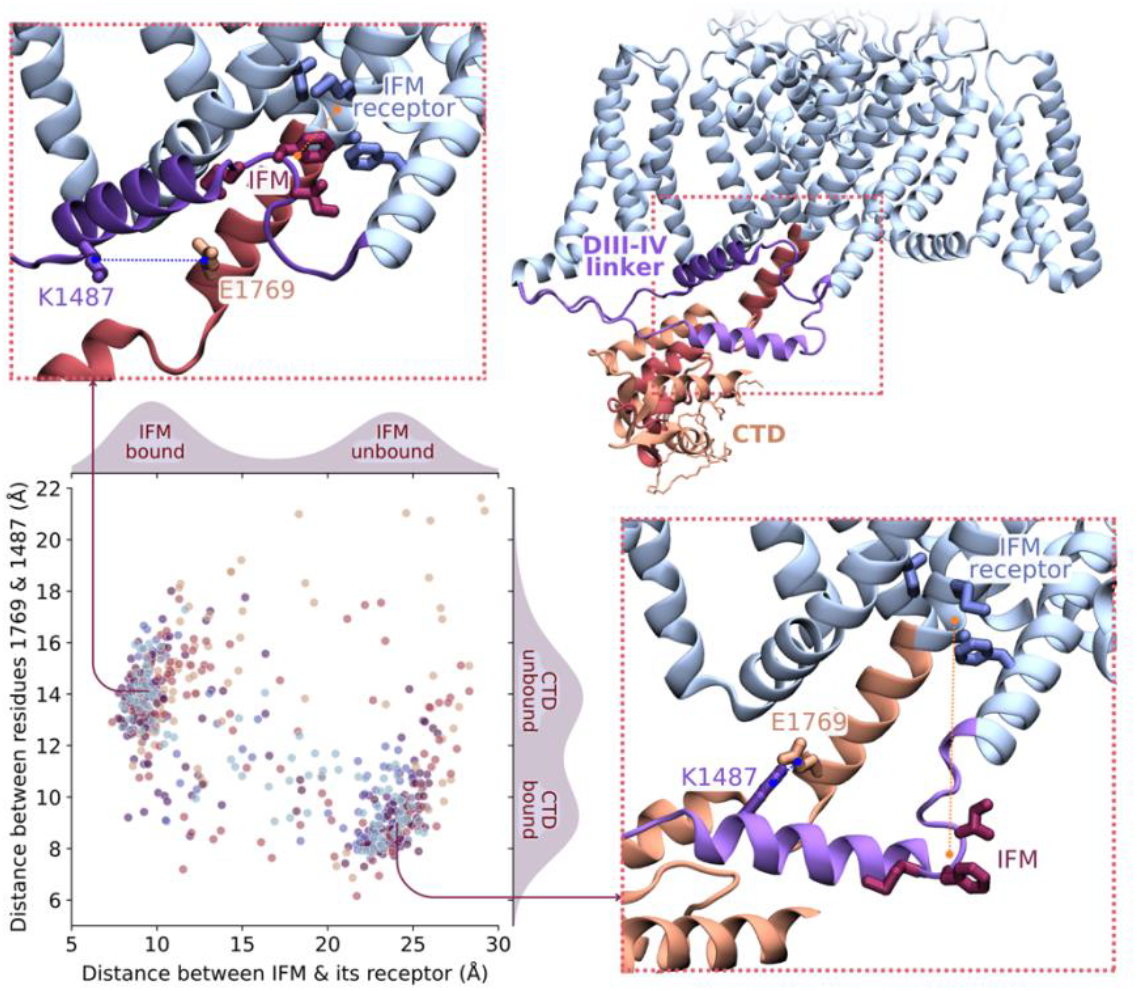
Conformation diversity in the DIII-IV linker and CTD region of Na_v_1.7. Distances were measured between IFM residues (I1472, F1473, M1474) and its binding site (F1460, A1647, I1754, M1757) in orange; and between residues E1769 on the CTD and L1487 on the DIII-IV linker in blue. Representative structures depicted from each of the clusters – top left representing IFM-bound and CTD-unbound state and bottom right representing the IFM-unbound and CTD-bound state.

## Discussion

Resolving the structures of alternative functional states of proteins, such as those in the VGCC superfamily, which undergo conformational changes in response to changes in membrane potential, remains a significant challenge. Therefore, we sought to explore whether AF2 could offer an alternative approach for determining the diverse structural conformations adopted by this protein family.

Our results demonstrate that AF2 can generate diverse conformations of VGCC proteins depending on the input sequence it is given. The AF2 ensembles we generated reveal a diverse array of structures that highlight the dynamic nature of the S4 helix in the VSDs. The vertical position of this helix in the ensembles often have multiple peaks that represent stable states with varying numbers of gating charge residues below the HCS. Some of the structures produced resemble existing cryo-EM structures, while others represent numerous transitional states, suggesting the plausibility of the models produced. Consistent secondary structural features are observed in our models that have been seen in previous structural and simulation studies including 3-10 helical regions in S4 and pi-bulges in S6. For the majority of VSD sequences folded, we see a bias for models to be generated in the activated S4-up state. In the cases of Ca_v_3.2 and Na_v_1.7, VSD4 produced the greatest conformational diversity both when modelling the VSD sequence only and when modelling the full protein sequence.

Despite capturing a diversity of plausible protein conformations there is evidence that these AF2 structures do not capture the full extent of the possible VSD conformational changes. Previous microsecond simulations of K_v_2.1 demonstrated that three gating charges traverse past the HCS during VSD deactivation in the presence of a large hyperpolarising membrane potential (Jensen et al., 2012) or biasing force (Delemotte et al., 2011). However, in the AF2-generated ensemble, we observe a maximum downward displacement of two gating charges from the activated position, but never in the downmost conformation seen in simulations. Thus, either the simulations have produced non-physiological conformations, or AF2 is limited in its ability to completely sample the conformational changes of the K_v_2.1 VSD. Conversely for HCN1, AF2 predictions show a much broader diversity in VSD states compared to what was previously observed with cryo-EM structures. These structures (Lee and MacKinnon, 2019) feature a break in the S4 helix when transitioning into a hyperpolarised state which was not present in the AF2 predictions of the HCN1 VSD only. This is likely due to absence of the pore domain and suggests that AF2 predictions of just the VSD may not represent physiologically relevant conformations.

Another interesting observation is that the conformational distributions of different functional regions are not always correlated in the way we expect. For example, since the activation of Na_v_ channel VSD-IV is coupled to its fast inactivation process, we would expect a more open pore structures to be biased towards the VSD-IV down state. However, this correlation is not evident in the AF2-generated tetrameric Na_v_1.7 ensembles. On the other hand, correlations between openness of the pore to VSD-I, II and III activation is difficult to assess given the lesser variety of conformations generated.

Across most AF2 predictions, we found that reducing the depth of the MSA input leads to a greater range of conformations being produced, more of which were further from the activated VSD state. The majority of models generated had a TM score of >0.9 compared to reference cryo-EM structures. This means we can generate more possible structures of the deactivated S4-down configuration of the S4 helix. However, insight into the MSA is important for understanding why this is the case and potentially guide us in ways to further manipulate the MSA for generating greater conformational diversity. In this study we also did not consider the use of templates to guide AF2 predictions which was previously shown to increase the number of conformations produced and bias them towards certain states (Ngo et al., 2024).

Here we see that generating a large number of AF2-predicted structures can provide greater insight than a single model. Although tempting to assume that greater frequency of one structure equates to greater likelihood or stability of that conformation, this may be beyond the scope of AF2 and needs to be verified for specific cases. Similarly, we cannot assume that a VSD predicted with diverse conformations has an S4 helix that is more mobile than a VSD that is only predicted in the activated state. It is likely the case that AF2-generated VSD structures are inherently biased towards the activated state due to overrepresentation of such structures in the PDB. Similarly, predictions of the full Na_v_1.7 sequence show a bias for the asymmetric closed pore state, which is a common theme across Na_v_ channel cryo-EM structures. Previous findings indicated that AF2 predictions of G-protein coupled receptors tended to more accurately produce inactive conformations likely due to the larger number of inactive structures in the PDB (Heo and Feig, 2022). Therefore, a shallower MSA reduces the evolutionary constraints on the prediction and reduce bias towards the overrepresented structures, as seen in previous work (del Alamo et al., 2022).

It has been shown that AF2 cannot be used broadly to predict the effect of a single mutation on protein stability (Pak et al., 2023). However, in certain cases of mutations in less flexible regions, AF2 has been able to identify disruption of the local structure (McBride et al., 2023). It is interesting to consider whether mutational effects can be uncovered by generating an ensemble of models. For example, could mutations that have been experimentally shown to shift the voltage dependence of activation of Na_v_Ab (McCord et al., 2020) lead to a bias in the conformational distributions of VSD models generated by AF2?

The ability to rapidly generate diverse structural states has several applications. First, plausible AF2-generated models in alternate conformations can be used to provide or reinforce mechanistic insight into channel behaviour, highlighting how this differs across the superfamily. Second, these models can serve as starting structures in simulation studies in which the structural models can be further refined or connected to yield trajectories between functional end states. The models can also be used to help define reaction coordinates used to guide further simulations of structural transitions. Third, they can help to address how different factors influence VSD or pore conformations. For example, with a range of naturally derived toxins and novel therapeutics that target specific states of the VSDs, and having structural details of the different states would be useful a useful basis for drug design. The tests employed in this work help assess the capabilities and limitations of AF2 structural modelling – including the extent of conformational heterogeneity that can be generated and the influence of different sequence inputs, as well as providing a wealth of information about the conformational diversity of VG cation channels.

## Methods

### Generation of AF2 models

Sequences for 36 VSD only constructs were extracted from full sequences of 12 different VG channels based on InterPro annotations in the voltage-dependent channel domain superfamily (IPR027359). A representative channel was chosen from each of the main ion channel families, which included Ca_v_1.1, Ca_v_2.2, Ca_v_3.2, Na_v_1.7, Na_v_Ab, rat K_v_1.2, rat K_v_2.1, K_v_3.1, K_v_4.2, K_v_5.1, K_v_6.1, K_v_7.1, K_v_8.1, K_v_9.1, rat K_v_10.1 (EAG1), K_v_11.1 (ERG), K_v_12.1 (ELK), TPC1, HCN1 and CatSper1.

ColabFold Version 1.5.2 (Mirdita et al., 2022) was used to generate MSAs and produce a total of 600 structures for each VSD. Five models were folded per five random seeds and repeated four times to predict 100 models per MSA depth condition (Fig. 1D). MSA depth was reduced five times by changing the –max-msa-seq parameter to 256, 128, 64, 32 and 16, without changing –max-msa-seq-extra (default at 5120). Number of recycles was set at 1 to minimise refinements that could lead to more homogenous structures. The –use-dropout flag was also applied to increase uncertainty in the predictions.

### Analysis of models

Sequence alignment of the VSD sequences was performed using ClustalW (Sievers et al., 2011). From this alignment, the residue positions of the conserved HCS and four centremost gating charge residues were identified for each VSD (relative to F56, and R99, R102, R105 and R108 in Na_v_Ab).

VSD conformational heterogeneity was evaluated by measuring the degree of change in the S4 helix of each model. Each model was aligned, using UCSF Chimera Structure Comparison MatchMaker (Pettersen et al., 2004), to the Na_v_Ab VSD that was pre-oriented in plane of the membrane. MDanalysis (Michaud-Agrawal et al., 2011) was used to measure the z-position of centre of mass (z_COM_) of the four gating charges, and subtracted from the z_COM_ of the HSC residue, in order to determine the relative z-displacement. The same procedure was used to measure the conformations of each of the four VSDs in the full tetrameric Ca_v_ and Na_v_ models. VSD conformations were differentiated into the number of gating charges below the HCS. This was determined by measuring the z_COM_ of each gating charge and counting the number located below the z_COM_ of the HCS. VMD (Visual Molecular Dynamics) (Humphrey et al., 1996) was used for visualisation of models and generation of structural images.

TM score (Zhang and Skolnick, 2005) analysis was performed to compare AF2 models to existing cryo-EM structures of Ca_v_3.2 -9AYG (Huang et al., 2024), Ca_v_2.2 - 7MIX (Gao et al., 2021) and Na_v_1.7 - 6J8G (Shen et al., 2019), 6NT4 (Clairfeuille et al., 2019) using the US-align standalone program (Zhang et al., 2022) with sequence dependent alignment and normalised to the length of the reference structure.

Ca_v_3.2 and Na_v_1.7 pseudo-tetrameric models were aligned to a membrane orientated structure of using Chimera. Pore conformations were then assessed by measuring the COM distance between S6 gating residues on opposite domains (DI-DIII distance and DII-DIV distance) using MDanalysis. Residues used for COM calculations for each of the domains were L415 and A419 on DI, V1011 and V1015 on DII, V1551 and V1555 on DIII, and V1856 and M1860 on DIV for Ca_v_3.2; and L398 and A402 on DI, L964 and L968 on DII, I1453 and I1457 on DIII, and I1756 and L1760 on DIV for Na_v_1.7. Distances were also measured for intracellular regions of Na_v_1.7, between IFM residues (I1472, F1473, M1474) and its binding site (F1460, I1754, M1757, A1647) and between residues E1769 on the CTD and L1487 on the DIII-IV linker.

Prevalence of 3-10 and pi helical regions in the S4 and S6 helices were measured using MDTraj (McGibbon et al., 2015) compute_dssp to identify the secondary structure across all the models. The z_COM_ of each residue was measured and plotted against the secondary structure.

Fenestration bottleneck analysis for Na_v_1.7 was conducted using Caver 3.0 (Chovancova et al., 2012), after adding hydrogens to the protein. The following parameters were used for tunnel calculation: probe radius of 0.8 Å, shell radius of 15 Å, shell depth of 15 Å, and origin set at the centre of mass of four residues at the midpoint of each S6 helix.

## Supplementary Figures

**Fig. S1.**
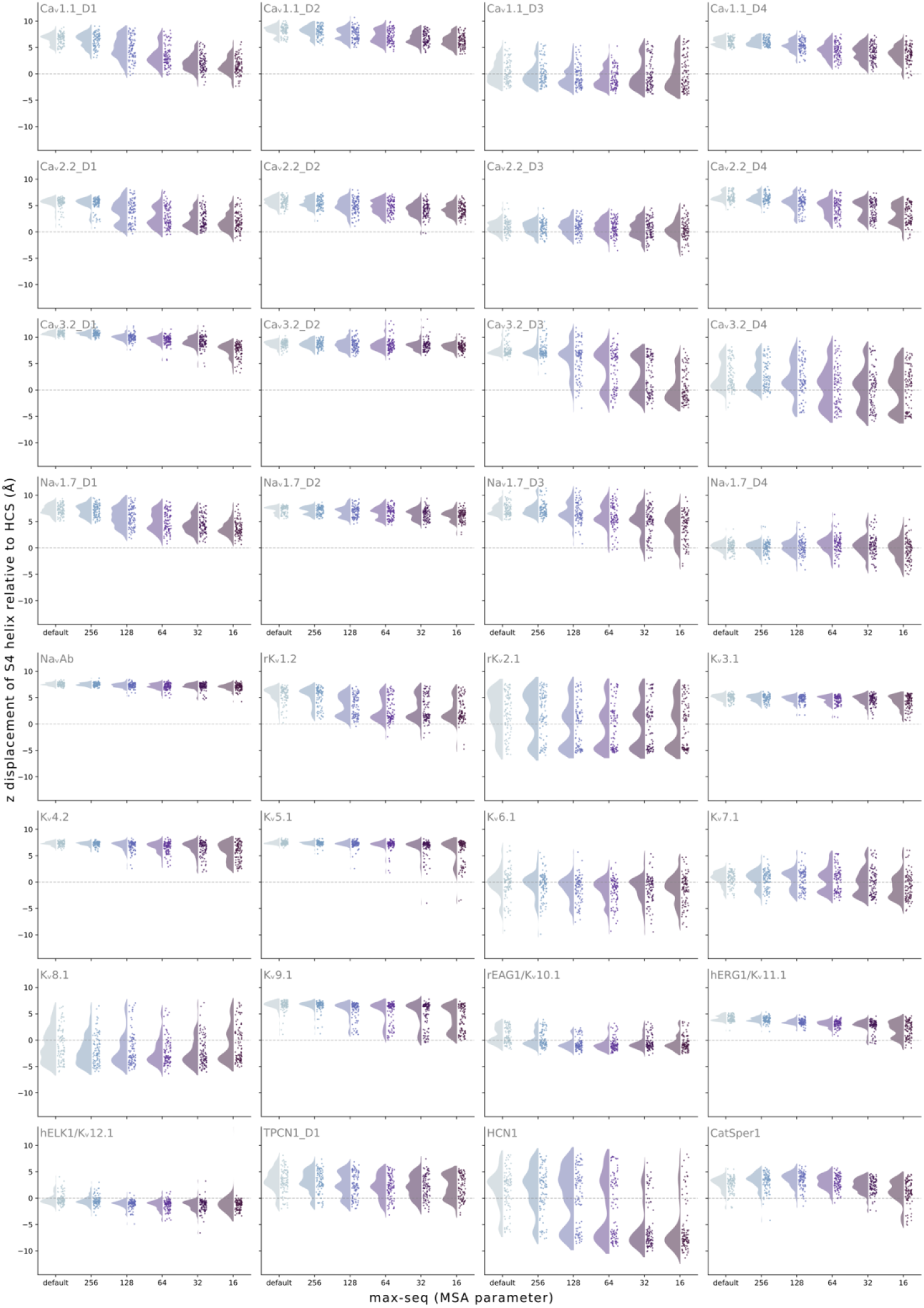
All conformational distributions of VSD only AF2-generated ensembles (shown in Figure 2), grouped by the max-msa-seq parameter used for predictions.

**Figure S2.**
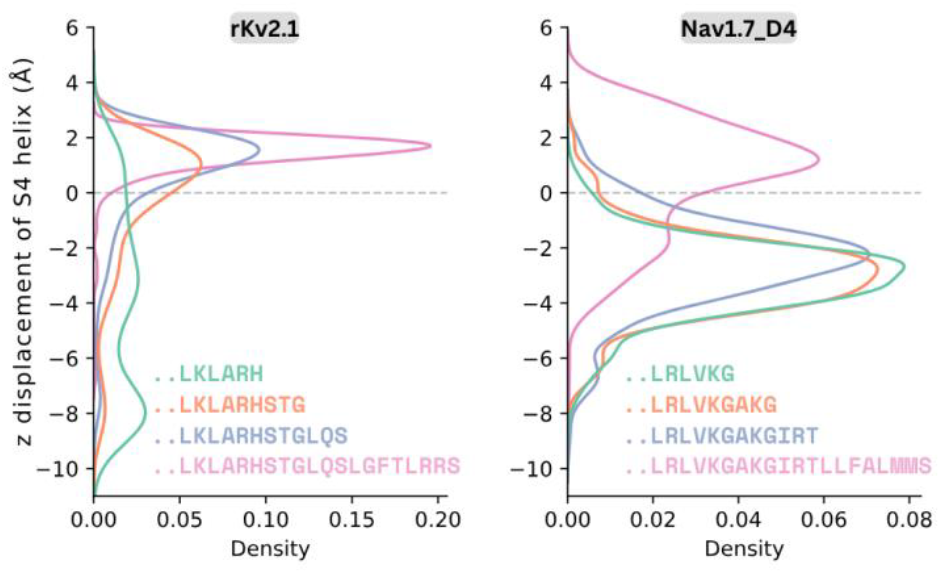
Addition of residues to the end of the VSD input sequence results in biased conformational distributions towards the activated VSD state for both the K_v_2.1 VSD and Na_v_1.7-VSD4.

**Figure S3.**
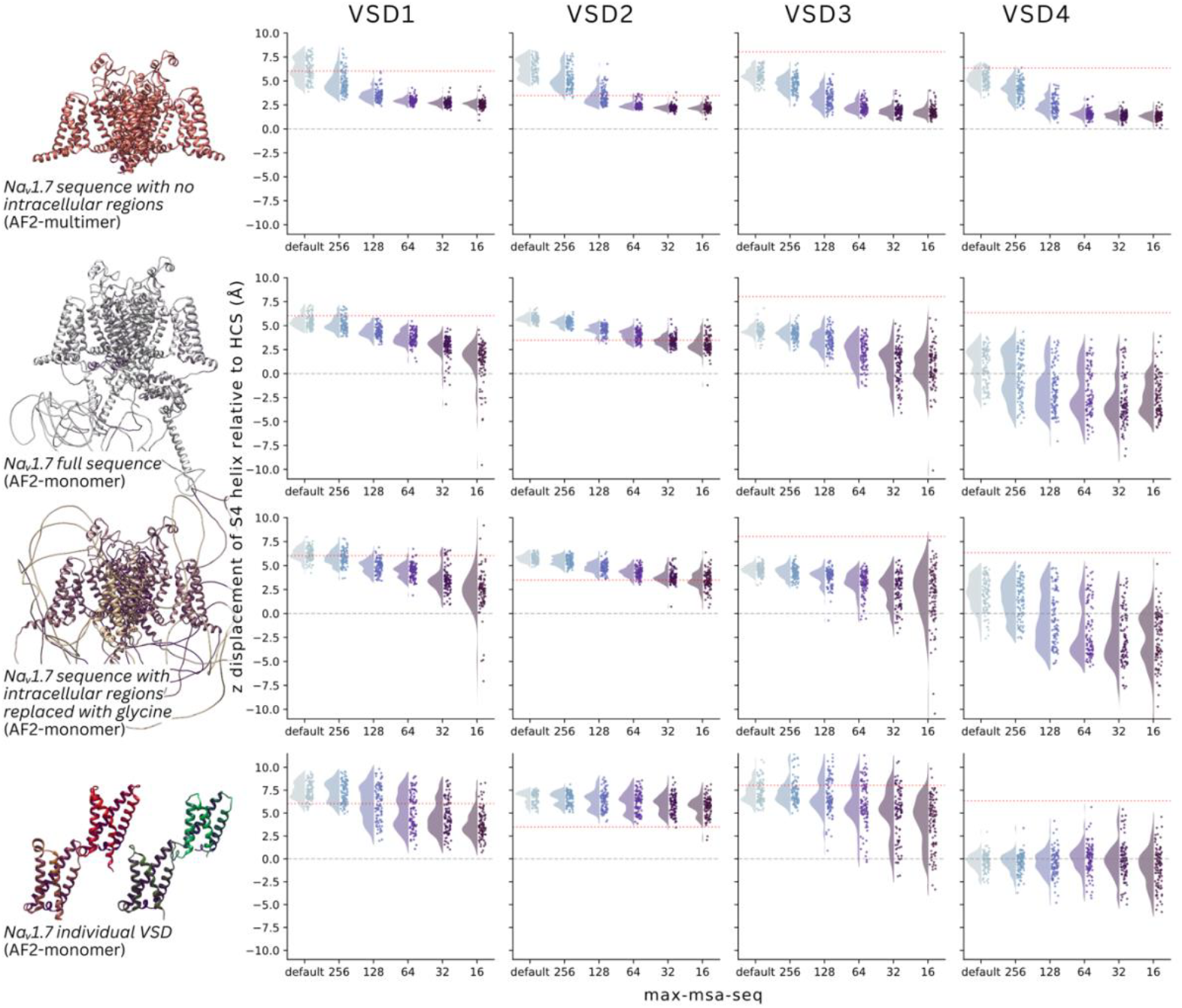
VSD conformational diversity in the models of Na_v_1.7 varies depending on the input sequence and the use of AF2 monomeric or multimeric version. The distributions show the z displacement of the S4 helix across 100 AF2 models for each of the max-msa-seq parameters. The four columns show each of the four VSDs. The rows show the different input sequence and AF2 version used, with a representative snapshot on the left. The red line denotes the VSD conformations found in the Na_v_1.7 cryo-EM structure, 6J8G.

**Figure S4.**
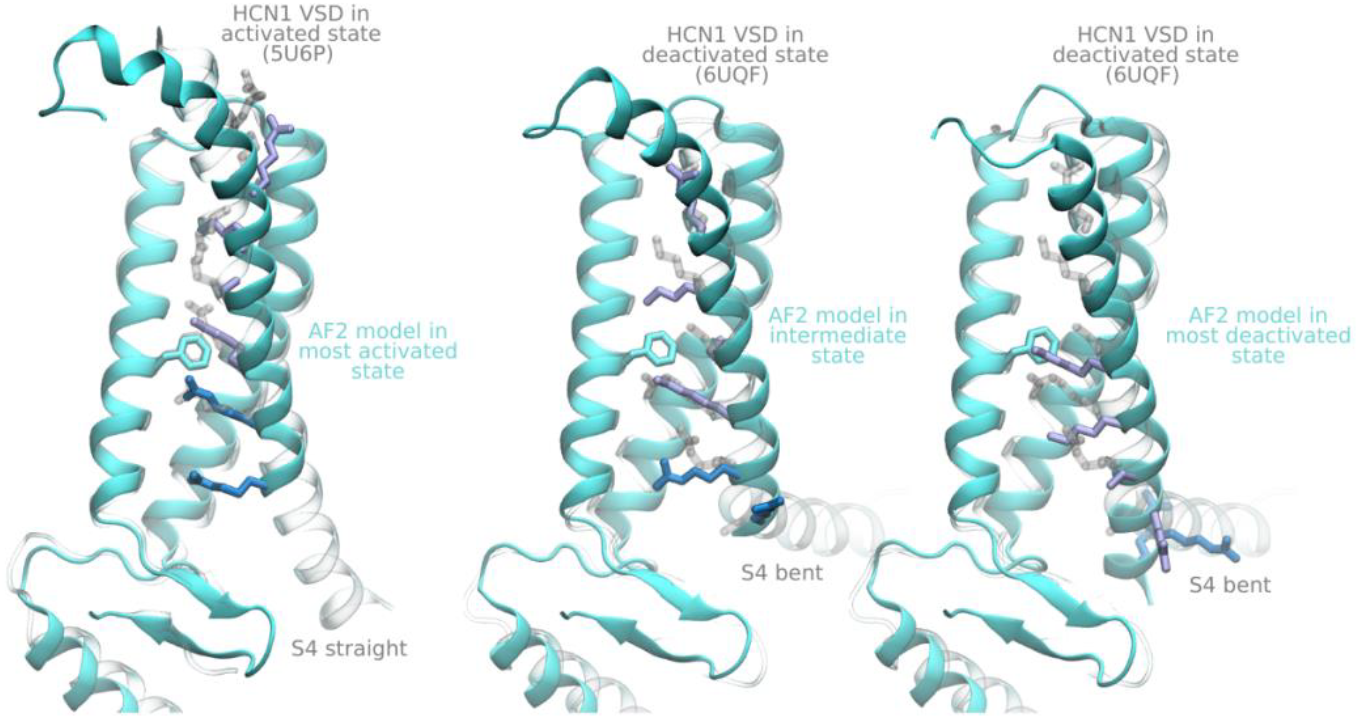
Structural alignment comparing AF2 models of the HCN1-VSD (cyan) to cryo-EM structures resolved in VSD-activated (left, transparent gray) and VSD-deactivated (middle, right, transparent gray) conformations, PDB IDs 5U6P and 6UQF respectively.

**Figure S5.**
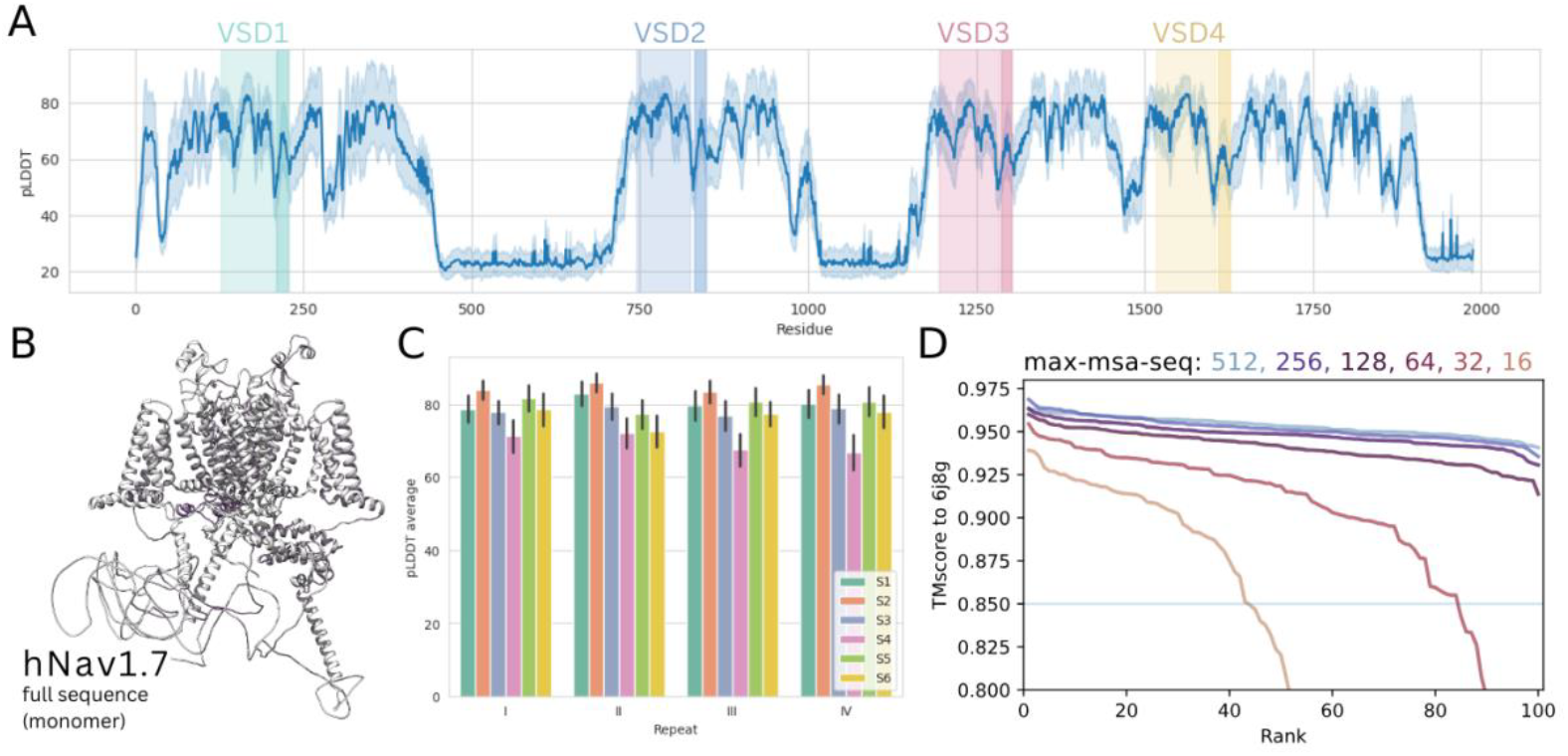
Analysis of AF2 predictions for the full Na_v_1.7 sequence. (A) pLDDT across all 100 Na_v_1.7 AF2 predicted models using default MSA parameters. (B) Representative snapshot of Na_v_1.7 model produced by AF2. (C) pLDDT values averaged across each of the 24 transmembrane helices. (D) TM score of Na_v_1.7 models to resolved cryo-EM structure 6J8G ordered by rank and coloured by MSA depth restriction parameter.

